# Addressing Heterogeneity in direct analysis of Extracellular Vesicles and analogues using Membrane-Sensing Peptides as Pan-Affinity Probes

**DOI:** 10.1101/2023.12.20.572525

**Authors:** Alessandro Gori, Roberto Frigerio, Paola Gagni, Jacopo Burrello, Stefano Panella, Andrea Raimondi, Greta Bergamaschi, Giulia Lodigiani, Miriam Romano, Andrea Zendrini, Annalisa Radeghieri, Lucio Barile, Marina Cretich

## Abstract

Extracellular vesicles (EVs), crucial mediators of cell-to-cell communication, hold immense potential for diagnostic applications due to their ability to enrich protein biomarkers in body fluids. However, challenges in isolating EVs from complex biological specimens hinder their widespread use. In this frame, integrated isolation-and-analysis workflows are the go-to strategy, most of which see the prevalence of immunoaffinity methods. Yet, the high heterogeneity of EVs poses challenges, as proposed ubiquitous markers are less homogenously prevalent than believed, raising concerns about the reliability of downstream biomarker discovery programs. This issue extends to the burgeoning field of engineered EV-mimetics and bio-nanoparticles, where conventional immune-affinity methods may lack applicability. Addressing these challenges, we introduce the use Membrane Sensing Peptides (MSP) as “universal” affinity ligands for both EVs and EV-analogues. Employing a streamlined process integrating on-bead capture and vesicle phenotyping through Single Molecule Array (SiMoA) technology, we showcase the application of MSP ligands in the integrated analysis of circulating EVs in blood derivatives, eliminating the need for prior EV isolation. Demonstrating the possible clinical translation of MSP technology, we directly detect an EV-associated epitope signature in serum and plasma samples, demonstrating its potential for distinguishing patients with myocardial infarction versus stable angina. At last, notably, MSP exhibits a unique capability to enable the analysis of tetraspanin-lacking Red Blood Cell derived EVs (RBC-EVs). Overall, unlike traditional antibody-based methods, MSP probes work agnostically, overcoming limitations associated with surface protein abundance or scarcity. This highlights the potential of MSP in advancing EV analysis for clinical diagnostics and beyond. Of note, this represents also the first-ever peptide-based application in SiMoA technology.

## Introduction

Extracellular vesicles (EVs) are nanosized, membrane-bound particles released by cells into the extracellular space^1^ and are known to play essential roles in cell-to-cell communication^2, 3^. EVs are arising unparalleled expectations in the diagnostic field, given their capacity to enrich potential protein biomarkers which otherwise constitute only a very small portion of the total proteome of body fluids (<0.01%) ^4, 5, 6, 7, 8^. The putative diagnostic power of EVs is particularly compelling for those biological specimens rich in EVs, including blood and urine, which allow for a non-ivasive sample collection with respect to tissue biopsies. However, despite considerable efforts, EVs isolation from complex body fluids remains arduous, time consuming and difficult to standardize. Combined with a growing body of evidence that pre-analytic steps in EVs sample preparation can bias the downstream analysis, multistep protocols for EVs analysis remains unsuitable for large biobanks screening and even less applicable in routinary diagnostic settings. As such, arguably, only integrated isolation-and-analysis workflows could enable the translation from research settings to the real usage of EV-associated biomarkers into clinical viable practices, with microfluidics ^9, 10, 11, 12^ and bead-based systems ^13, 14^, ^15^ appearing as the most promising methods.

In these systems, immunoaffinity often remains a standard to isolate and analyze EVs, with typically used targets for isolation including tetraspanins (CD9, CD81, CD63) and other EV-surface proteins such as EpCAM or EGFR. However, with the surge of sorting methods and single-vesicle analysis techniques, the high heterogeneity of EVs is clearly emerging: several markers proposed to be ubiquitous are less prevalent than believed, and multiple biomarkers patterns concur in single vesicles but only in small sub-fractions ^16,17, 18^. Thus, affinity purification based on specific EV surface proteins, if no prior knowledge is generated by other techniques (e.g. proteomics), can lead to missing information and be misleading, undermining the reliability of downstream biomarker discovery programs. It is paradigmatic that a recent work reported on the loss of up to the 80% of EVs with diagnostic potential when affinity isolation by single tetraspanin protein is used, and a loss of 36–47% when a tetraspanin cocktail is employed ^19^.

This issue reverberate also on the exponentially growing field of engineered^20^, synthetic^21^ or hybrid EV-mimetics^22, 23^, and other bacterial ^24, 25^ or plant derived bio-nanoparticles^26^, which are under extensive investigation as the next generation nanodelivery systems in the therapeutic arena. Indeed, their successful translation depends, among other factors, also on the availability of high-sensitivity characterization methods to be applied into robust QC protocols, and to assess biodistribution parameters including their concentration and clearance in biofluids. In this frame, conventional immune-affinity methods, typically used for “natural” EVs, may not be readily applicable due to poor knowledge or scarce abundance of surface markers, or unavailability of validated antibodies Based on these rationales, we have previously introduced^27^ the use of Membrane Sensing Peptides (MSP) as a class of “universal” affinity ligands for nanosized lipid particles, including small EVs (sEVs, <200nm size range). In the context of EVs and their analogues or mimetics, rather than targeting a specific protein on the surface, MSP show specific affinity for the highly curved membrane, which can be considered a shared pan-vesicular epitope, making MSP probes working agnostically in regard to the relative abundance of surface proteins in different EVs sub-populations or EV-mimetics compositions (Figure 1). More specifically, curvature sensing is a result of lipid packing defects characteristic of highly tensioned membranes, which favor the membrane insertion and binding stabilization of amphipatic protein domains or peptides ^28, 29, 30, 31^.

**Figure 1:**
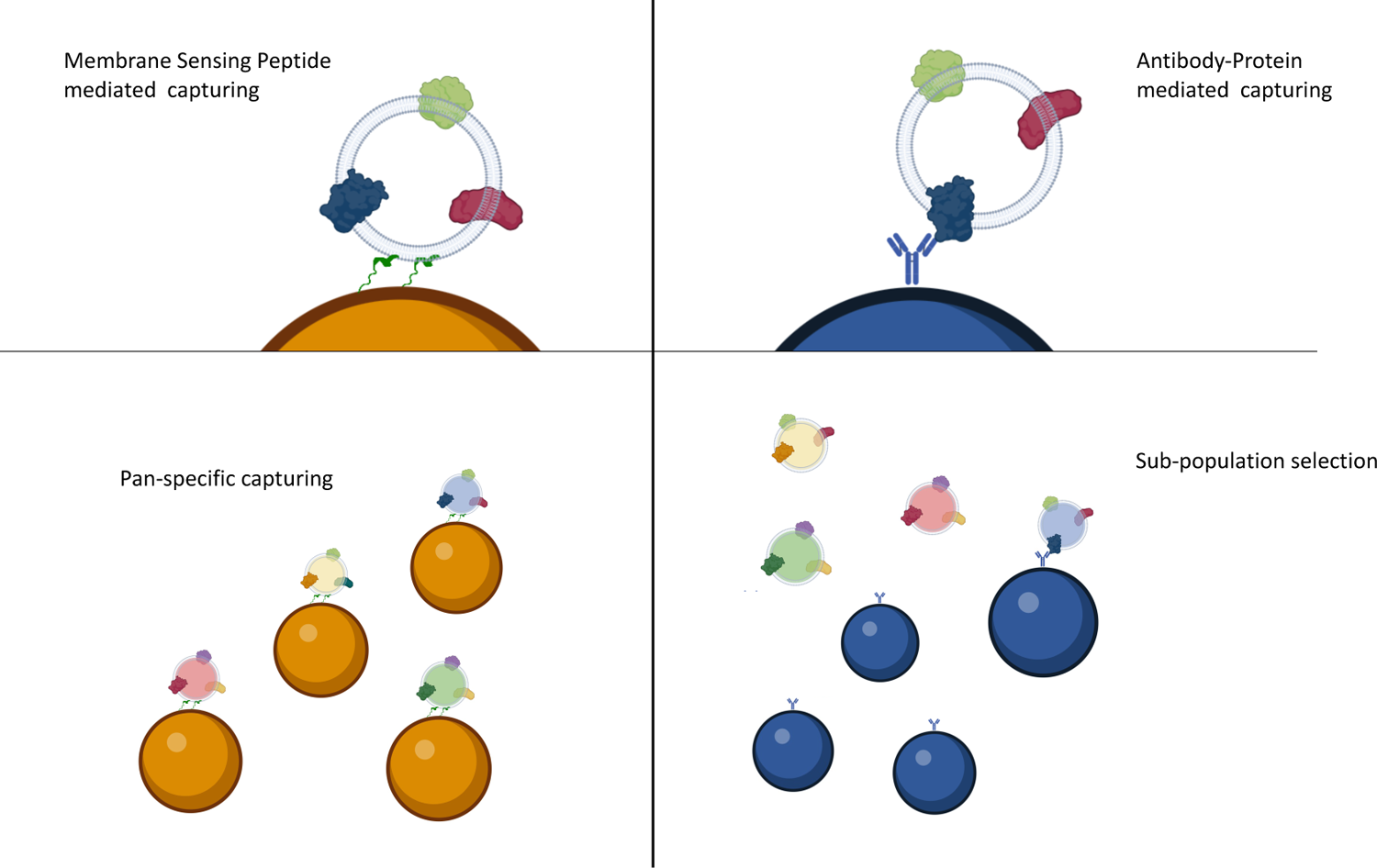
Membrane Sensing Peptides (MSP) – left panel - are able to capture small EVs or analogues based on their membrane physical traits, irrespectively from surface protein abundance, rendering a pan-specific capturing. On the contrary, antibody-mediated capturing – right panel – depends on the presence of specific protein epitopes on the EV surface, selecting only specific sub-populations

Here we present the use of MSP ligands for the integrated analysis of circulating EVs in blood derivatives (serum and plasma) without the need for prior EV isolation, and, to further demonstrate the versatile application of MSP probes in EV analogues analysis, we extend this approach to an emerging class of EVs, Red blood cell derived EVs (RBC-EVs) ^32^. Obtained by calcium ionophore stimulation, RBC-EVs show peculiar surface characteristics such as lack of canonical tetraspanins (CD81, CD9 and CD63) and phosphatidylserine enriched surface ^33^.

Specifically, we relied on a streamlined process that integrates on-bead capture and vesicle phenotyping through the Single Molecule Array (SiMoA)^34,35^ technology. The unparalleled sensitivity of this digital detection platform, integrated into the high-throughput Simoa® instruments, favoured its diffusion in clinical centers for monitoring of biomarkers in the field of neurological diseases, cancer and other chronic diseases. Recently, SiMoA significant clinical potential have been largely documented in EV analysis ^36, 37, 38, 39^.

Efficiency of MSP versus antibody-based capturing is shown, and the unique capability of MSP to enable analysis of tetraspanin-lacking RBC-EVs is reported. Finally, we demonstrate possible MSP translation in clinical settings by directly detecting an EV-associated epitope signature in a cohort of serum and plasma samples for stratification of patients with myocardial infarction *versus* stable angina. Of note, this represents also the first ever peptide-based assay in the SiMoA technology.

## Results and discussion

### MSP selectivity of EV binding in serum and plasma

Blood is a highly complex matrix that proved to be challenging for reproducible EV isolation and biomarker analysis ^40^. The combination of more than one method, for example density cushion and size exclusion chromatography, is often needed to efficiently purify EVs from other blood components, mostly including lipoproteins, which largely outnumber EVs ^41, 42, 43^. Yet, gold standard techniques for EV pre-isolation, or combination of orthogonal purifications, are impracticable for large cohort studies or routine diagnostics.

As such, the integration of EV isolation and analysis in a one-step streamlined protocol is an highly appealing strategy to pave the way to actual clinically compliant procedures. In view of their direct use in blood specimen without pre-isolation steps, we set to assess whether MSP probes meet the criteria of specificity of EV capturing with respect to common contaminants in blood derived specimen. Of note, MSP were recently proposed for EV isolation from cell conditioned medium by capture and release from modified agarose beads, rendering a fully competent sEV preparation protocol with performance superior to most of the current commercially available isolation kits ^44^. Here, we applied the same format of isolation to screen MSP capturing efficiency of EVs from serum and plasma, and to monitor the binding selectivity versus the most abundant blood “contaminants” in EVs analysis, including lipoproteins (Apolipoprotein A, Apolipoprotein B, Apolipoprotein E) and albumin. This was verified by a catch-and-release strategy (Figure 2-A, see Materials and Methods section), enabling both their characterization and the detection of possible co-isolated contaminants. Released EVs were fully characterized according to MISEV guidelines^1^ in terms of integrity, size and morphology by TEM (Figure 2-B, 2-C) and Nanoparticle Tracking Analysis (Figure 2-D), whereas Western Blot (Figure 2-E) was used to confirm the presence of typical EV-associated surface (CD63 and CD81) and luminal (TSG101 and Alix) markers. Overall, morphology, size distribution, and characteristic molecular markers of released sEVs were shown to be coherent with broadly accepted standards. The selectivity of MSP-driven EVs isolation was then tested in different blood pre-analytical conditions as they are well known to strongly influence downstream results ^45^. Immuno-dot blot analysis was used to assess possible co-isolation of albumin and of lipoproteins, via the use of anti-human serum albumin (HSA) and anti-Apolipoprotein A, B and E antibodies. Dot blot analysis for serum is reported in Figure 2-F, showing negligible signals for contaminants in the EVs released fraction. Similar results were obtained with plasma, independently from the pre-analytical conditions (Figure S1, Supplementary Information). Overall, we provided evidence that MSP probes were eligible candidats for integrated EV isolation-analysis in blood derived specimen, with minimal matrix and pre-analytical conditions interference.

**Figure 2:**
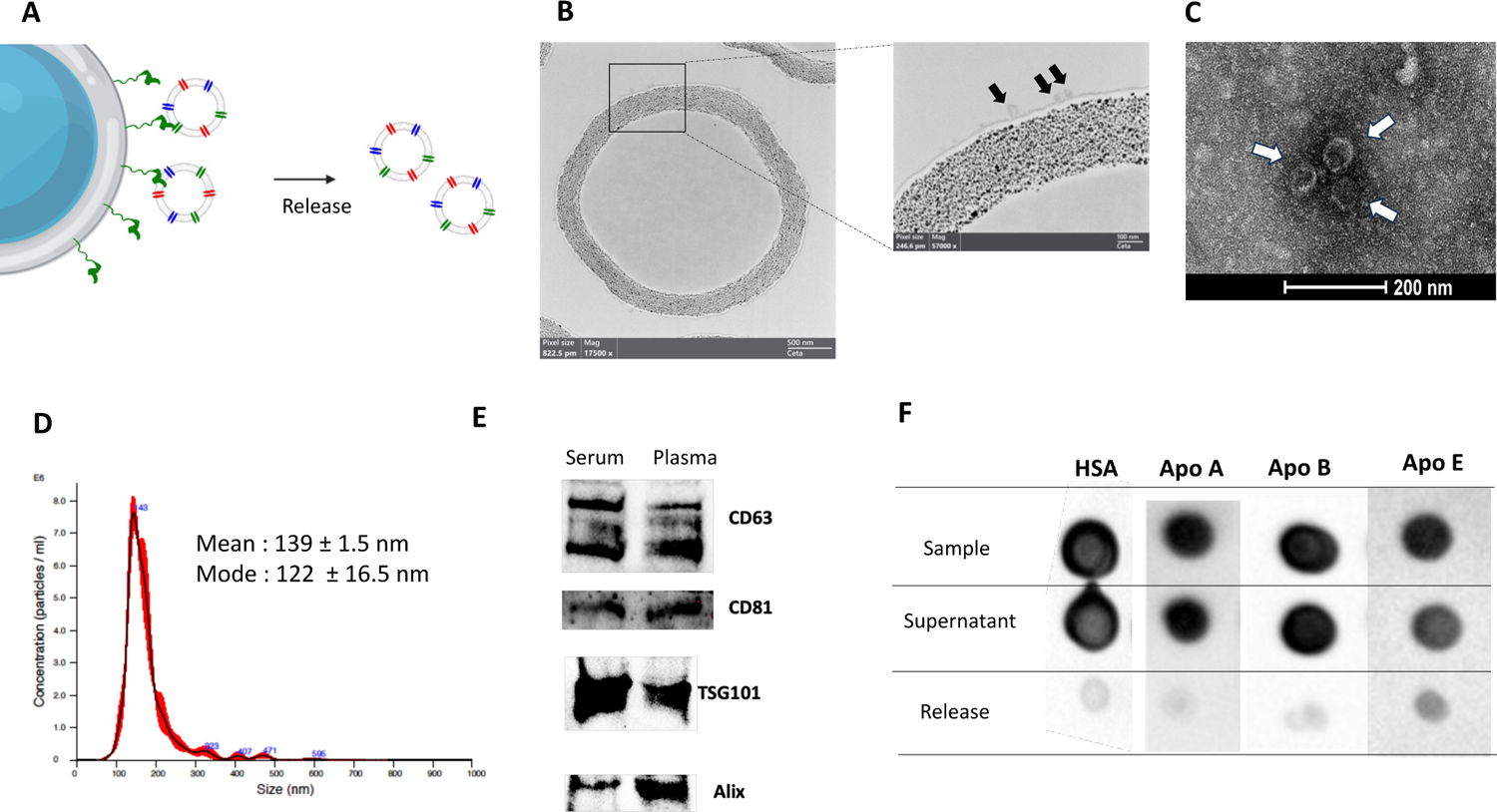
**A**) scheme of the capture and release strategy here applied to demonstrate efficient isolation of small EVs, their release in intact conditions and negligible presence of contaminants after separation. **B**) TEM image of agarose beads and surface captured EVs. **C**) TEM image of EVs captured from serum after release with 0.5M imidazole, EVs appear to be intact and present the typical cup-shape. **D**) NTA analysis of EVs captured from serum and released after imidazole treatment. Overestimation of EV diameter by NTA could be ascribed to instrument inability to detect EVs smaller than 70 nm. **E**) Western Blot detecting typical EV-associated surface (CD63 and CD81) and luminal (TSG101 and Alix) markers in the released particles. **F**) Immuno dot-blot analysis to check presence of common contaminants: Human Serum Albumin (HS) Apoliporoteins A, B and E (Apo A, Apo B, Apo E respectively) in the starting Sample (Serum), in the Supernatant after capturing and in the Released EV fraction. Negligible contaminant signals are detectable after EV isolation. Results form other pre-analytical conditions tested, plasma-EDTA, plasma-heparin and plasma-citrate are shown in Supplementary Information Figure S1.

### MSP capture returns representative EV markers abundance profile

Prior to application into real context scenarios, we more deeply investigated the representativeness of MSP binding propaedeutic to immunephenotyping, taking as a reference EVs with “canonical” surface marker profile. In other words, by testing three markers of consolidated use (CD9/CD63/CD81), this approach wanted to assess whether the outcome of analysis obtained by using MSP probes is reliable and results in no specific enrichment of some EVs subpopulations. Specifically, we set to verify that the EVs capture mediated by MSP returns the same level of tetraspanins (CD9/CD81/CD63) relative abundance with respect to that provided by the corresponding antibodies. This was performed directly on the analytical system of intended final use, namely the SiMoA Bead Technology, and directly into plasma from six healthy donors and in compliance with an integrated isolation-and-analysis protocols. SiMoA is a high-throughput and ultra-sensitive platform that we have previously explored for digital EV immune-phenotyping ^46^. As a reference, SiMoA microbeads were functionalized with a combination of antibodies directed against of CD9, CD63, CD81 tetraspanins (pan-tetraspanin beads, hereafter referred to as Tetra beads). These would serve as a proxy of global and unbiased capture beads for tetraspanins-positive EVs, regardless of the relative abundance among the three markers.

We then compared results obtained by alternatively using MSP beads *vs.* Tetra beads (Figure 3-A) for EVs capture, followed by surface immune-phenotyping to detect CD9/CD63/CD81 tetraspanin individually. Overall, an almost overlapping pattern of each tetraspanin relative abundance was observed either by using MSP beads (Figure 3-B, left panel) or Tetra beads (Figure3-B, right panel).

**Figure 3:**
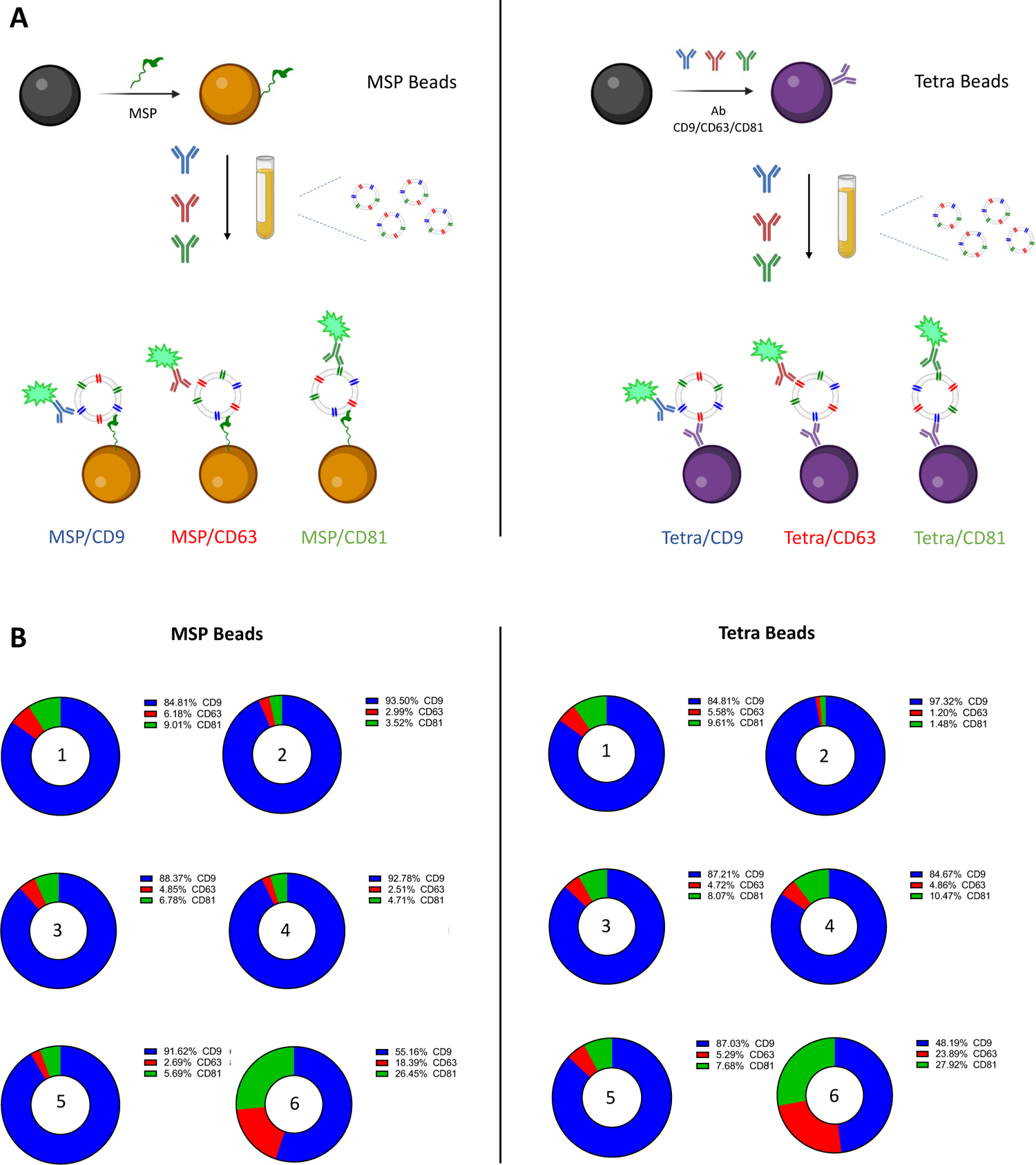
**A**) SiMoA Beads were modified by MSP (left panel) and by a combinantion of antibodied directed against CD9/CD63/CD81tetraspanin (right panel). Single tetraspanin immune-phenotyping of plasma EVs from 6 healthy donors was run in parallel on the two types of beads. **B**) For both settings and each sample, the total AEB (Average Enzyme per Bead) is calculated as the sum of single CD9/CD63/CD81 AEB detection. Single tetraspanin expression level is calculated as the AEB% over the total AEB. Both methods confirm the expected heterogeneity of EV samples with remarkable accordance of the two systems in terms of differential tetraspanin profiling.

This suggests that MSP binding does not result in a biased selection of EVs, nor that specific subtypes are enriched, and thus confirms their representativeness and reliability towards their perspective use in blind EV-associated biomarker discovery programs.

In these regards, it is worth noticing that, even if in this specific context (plasma EVs) a pan-tetraspanin capture is apparently well performing, it could not be the case for other EV samples where tetraspanins are poorly expressed and/or alternative abundant surface markers are not known.

Moreover, the relative abundance of CD9, CD63, CD81 highlighted the (expected) sample heterogeneity and the well known uneven distribution the three proteins ^16^ (Figure 3-B). To the best of our knowledge, the vast majority of published work in EV analysis after immune-based enrichment make use of individual anti-tetraspanin probes, without a prior pre-assessment of their relative abundance. This poses obvious concerns about the possible biases that can be introduced in downstream processes, especially when a sub-population is selected prior to biomarker analysis, thus re-inforcing the need for “universal” enrichment methods. All of this aside, MSP represent valuable alternatives to pan-tetraspanin capturing.

### Validation of EV immunophenotyping workflow by MSP in clinical settings

It was previously demonstrated that EVs analysis can reveal the very early stages of cell stress that precede cardiomyocytes death and the release of troponin, the biomarker in clinical use for the diagnosis of the acute coronary syndrome (ACS). More specifically, it was shown that significantly increased levels of EVs (monitoring of CD9/CD81/CD63 markers) and of vesicular CD42a, and CD62P antigens—endothelial and platelet-related antigens— were a distinctive signature in serum samples from patients experiencing ST-elevation myocardial infarction (STEMI), an acute coronary event that precedes myocardial injury ^47^.

In this frame, while replicating the full clinical validation was out of the scope of the current work, we still aimed at validating the use of MSP probes in this clinically relevant context, using an integrated isolation-and-analysis protocol that would match a viable workflow for direct EVs analysis from blood-derived specimen. Besides assessing the technical feasibility, it was crucial to determine whether MSP-based EVs epitope profiling would return the same diagnostic value obtained by gold standard platforms encompassing the use of antibodies as EV-binding probes, such as the one that was used in the previous study by Burrello et al ^47,48^. We therefore performed an MSP-SiMoA assay for a selected panel of EV-associated markers proposed to serve in STEMI diagnosis by probing both serum and plasma samples without any form of EVs pre-isolation or enrichment. We evaluated the expression levels of CD9/CD81/CD63 tetraspanins, as well as those of vesicular CD42a and CD62P antigens, and assessed their value in the stratification of patients experiencing STEMI (n=12) vs. those sympomatic with stable angina (SA, n=12), who were not undergoing an acute ischemic event. These groups were carefully matched in terms of age, sex, and cardiovascular profile, utilizing a cohort previously described in the study by Burrello et al. ^47^. Our findings revealed higher serum levels of all evaluated EV-associated markers in serum of STEMI patients compared to SA (Figure 4).

**Figure 4:**
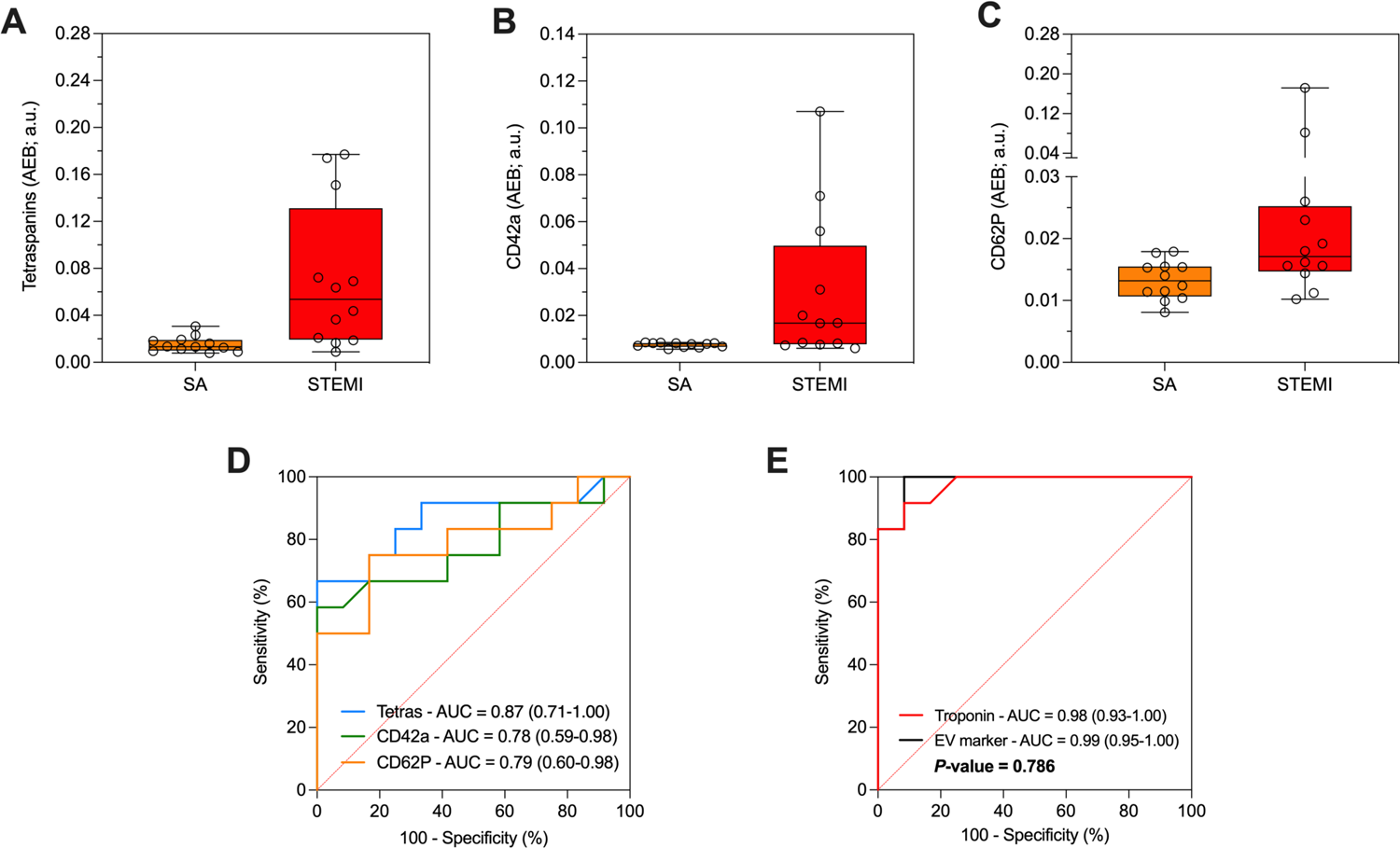
**A)** Expression of CD9/CD81/CD63, **B)** CD42a and **C**) CD62P in serum of patients with ST-segment elevation myocardial infarction (STEMI; red, n=12), stable angina (SA; orange, n=12). **D)** ROC curve analysis for individual EV markers CD9/CD81/CD63 (blue curve), CD42a (green curve) and CD62P (orange curve) **E)** Diagnostic performance of EV markers, compared with hs-troponin. ROC curve analysis of hs-troponin (red curve) compared to an aggregate EV marker including EV tetraspanin, CD62P, and CD42a; (black curve). The study was conducted with MSP modified beads in a customized SiMoA assay as described in the Materials and Methods Section.

The diagnostic performance of EV surface markers in discriminating STEMI patients and SA was assessed by ROC curve analyses (Figure 4-D). In the training cohort, ROC curves indicated a high sensitivity for these markers. An aggregate marker including the three EV parameters (EV tetraspanin, CD62P and CD42a levels) was compared with classical high sensitive troponin assay (hs-troponin) (Figure 4-E). AUC confirmed excellent diagnostic performances of the aggragate marker (0.99; 95% CI: 0.95-1.00), comparable with hs-troponin alone (0.98; CI: 0.93-1.000). Overall, we thus confirmed that using the proposed EV markers in a MSP-SiMoA assay returned diagnostic performances not inferior to hs-troponin (p=0.786). To further validate the reliability of our approach in different starting materials (plasma versus serum), a correlation analysis in patients with STEMI or SA (Figure 5) was performed.

**Figure 5:**
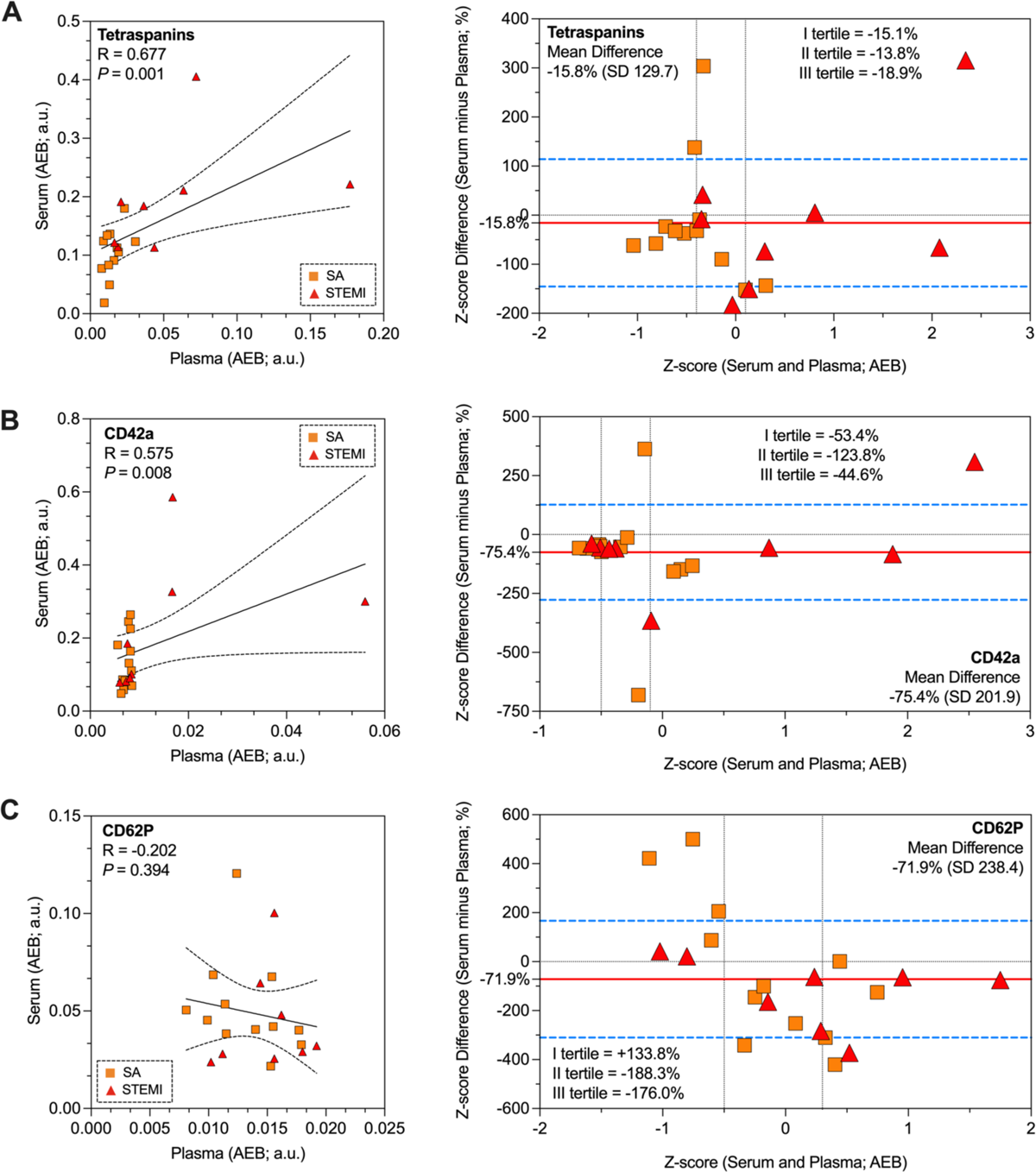
Expression levels in serum and plasma were correlated by Pearson’s R test in patients with STEMI or SA (n=12; left column). Bland Altman plots evaluating serum and plasma expression of tetraspanins (**A**), CD42a (**B**) and CD62P (**C**) after normalization by Z-score (n=12; right column). Difference between serum and plasma expression levels is reported on Y-axis; mean expression in serum and plasma is reported on X-axis, for each EV marker. The red line indicates mean percentage under-estimation of expression levels in serum compared to matched plasma samples, together with 95% confidence interval (blue dotted lines); tertiles of expression in serum and plasma are marked on the X-axis, together with the mean difference of serum minus plasma in each of them.

We observed a significant correlation for tetraspanins, and CD42a, but not for CD62P, in serum and plasma samples (with R values ranging between 0.499 and 0.677). Additionally, Bland Altman plots analysis demonstrated a consistent underestimation of the levels of expression of EV-associated epitopes in serum compared to plasma (Figure 5). No proportional or magnitude-dependent biases were observed for tetraspanins and CD42a. However, CD62P appeared to be overestimated in serum compared to plasma for lower expression levels (I tertile = +134%) and underestimated for higher levels (II tertile = −188%; III tertile = −176%). This made it challenging to directly compare CD62P levels in the two biofluids. In this sense, we suggest that marker of activated platelets could be biased by pre-analytical factors, due to uncontrolled platelet-activation during plasma collection.

### MSP in tetraspanin-lacking EVs samples

To further demonstrate surface-protein independent capturing of EVs as one of the key advantages of the MSP technology, we selected EVs derived from induced Red Blood Cells (RBC-EV). These vesicles are under investigation as candidates for drug delivery and other translational applications due to their high safety profile and minimal risk of horizontal gene transfer ^33^. In order to increase EV production, vesiculation can be in vitro induced by calcium ionophore, which also triggers phosphatidylserine flipping from the inner to the outer membrane leaflet while boosting EV release. RBCs lack the endolysosomal system hence they generate EVs only by plasma membrane budding (i.e. ectosomes), not expressing the canonical exosome tetraspanins (CD81, CD63, CD9) (See Supplementary Information Figure S2) but highly enriched in erythrocyte specific Band 3 anion transport protein (Band 3), and other EV markers like Flotillin-1, Alix, Annexin XI, and the lysosomal associated membrane protein 1 (LAMP1) ^49,50^.

As such, conventional EVs immune-affinity tools designed on tetraspanins are likely of low performance in the characterization and analysis of RBC-EVs. To prove this assumption, MSP beads and Tetra beads were again compared in RBC-EVs analysis on SiMoA (Figure 6-A) using anti-Band 3 as detection antibody and serial dilutions of an RBC-EV sample. Results are reported in Figure 6-B. MSP beads, differently from Tetra beads, are clearly able to specifically capture RBC-EVs down to 10^8^ vesicle/mL particle concentration (NTA determined)

**Figure 6:**
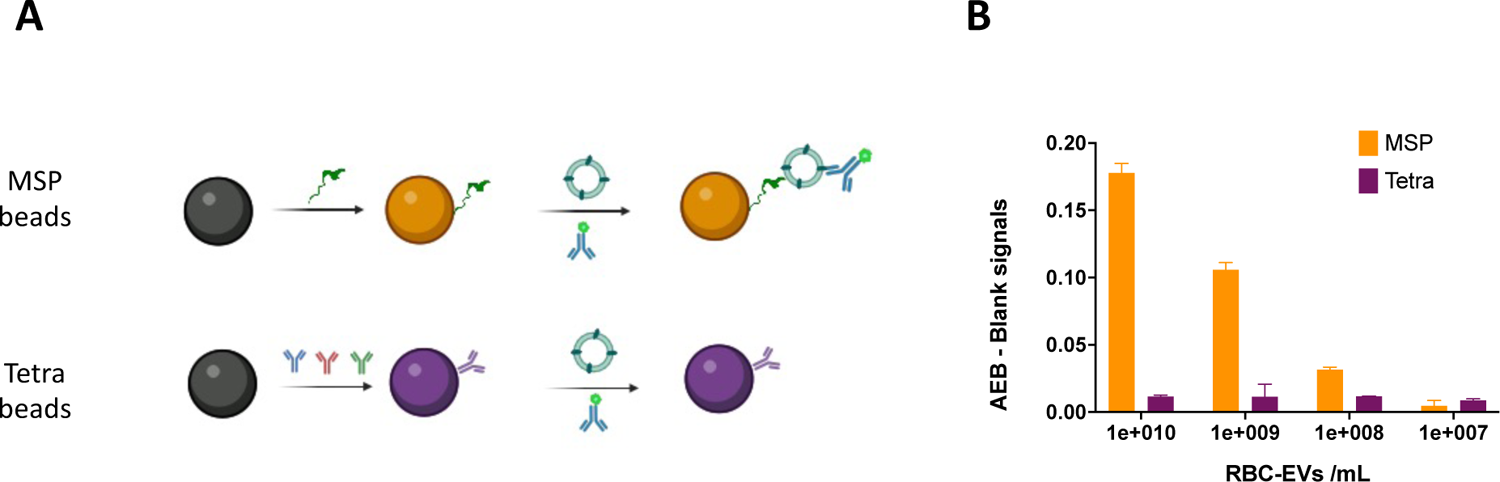
**A**) SiMoA beads were mofied by MSP and by a combination of CD9/CD63/CD81 antibodies (Tetra) and used to capture serial dilutions of RBC-EV using anti-Band 3 antibody for detection. **B)** Detection signals reported as Average Enzyme per Bead (AEB) subtracted of the corresponding blank signals are reported showing capacity of MSP beads to capture RBC-EVs down to 10^8^ vesicles/mL

## Conclusions

We have reported on a new tool for integrated capture and analysis of blood derived sEVs. The workflow is based on MSP, a new class of affinity ligands able to enrich small vesicles on the basis of specific membrane biophysical traits, thus avoiding pre-selection of EV sub-populations introduced by the use of antibodies. Our system is a new, flexible and blind biomarker discovery platform suitable for sampling sEVs directly from complex biological matrices like serum and plasma, with minimal carry over of contaminants and allowing subsequent EV epitope analysis in a one-step process. MSP presents several advantages when compared to systems relying on immune-capturing as the first step of analysis: *i)* MSP do not introduce biases in terms of sub-population selection upstream EV biomarker analysis; *ii)* MSP can be used to enrich EVs when there is poor knowledge of the most abundant EV surface markers or there are no evidence of highly expressed proteins as proxy of general capturing *iii)* MSP are not species-specific and can be used for samples from animal or plant species against which no validated antibodies exist; *iv)* MSP shares typical advantages of peptides over antibodies such as cost-effectiveness, longer shelf-life and no batch-to-batch variability.

Our MSP technology underwent demonstration for compliance with specificity criteria in a catch and release protocol of EV isolation from plasma and serum in different pre-analytical conditions. Then it was implemented, as a first example of peptide-based SiMoA assay, into a streamlined capture and analysis workflow, demonstrating to be a viable alternative to pan-tetraspanin capturing and especially useful with samples lacking of CD9/CD63/CD81 such as RBC-EVs, setting itself at the forefront of technologies for manipulation and analysis of next-generation bio-nanoparticles.

MSP capturing was validated in a clinically relevant context based on previously published evidence highlighting the significance of surface EV antigens in the context of cardiovascular diseases ^48^. The potential diagnostic performance of EV markers for STEMI showed non-inferiority to gold standard assay based on hs-troponin. Due to ease of use and adaptability, we foresee the wide adoption of MSP, and their integration into various isolation and analytical platforms, including the Simoa® instrumentation. This technology serves as a versatile toolkit for small extracellular vesicle enrichment and analysis, aiming for seamless incorporation into clinical automated routines.

In these scenarios, we expect the use of MSP beads within open EV biomarker discovery platforms to selectively enrich small extracellular vesicles, even from complex samples such as serum and plasma. This pan-selective enrichment process would precede the subsequent immune-phenotyping of putative biomarkers of interest, protecting the biomarker discovery process from bias introduced by the use of antibodies in the capturing step.

## Acknowledgements

We thank Prof. Paolo Bergese for the stimulating scientific discussions, unconditional support and friendship. Work was partially funded by the European Union through Horizon 2020 research and innovation program under grant agreement No. 951768 (project MARVEL) and by Next Generation EU trough MUR-PRIN 2022 project EV-PRINT (2022CS9H53), and MUR-PRIN 2022 project 2022BLLAEX

## Materials and Methods

### EV Isolation via agarose beads and characterization

High Cobalt density agarose resins from Agarose Bead Technologies (ABT) were conjugated with MSP-H6 peptide (6XHis-RPPGFSPFR-RPPGFSPFR) as described in Benayas et al ^44^. For blood derived specimen EVs isolation, 0.1 mL of MSP-beads suspension was added to 50uL of sample, serum or plasma EDTA, plasma heparin or plasma citrate diluted 1:10 in PBS, to a final volume of 500uL and incubated on RotoFlex for 1 hour at room temperature. Using a magnetic stand, supernatant was recovered, then beads were washed three times with 0.5mL PBS. EVs release was performed adding 100uL of Imidazole solution 0.5M in PBS for 15min under shaking, at room temperature and EV suspension recovered usig a magnetic stand.

### Dot and western blot analysis of released fraction

For analysis of contaminants, 3uL of pure sample was dropped off on nitrocellulose membrane (Protran BA 85 Nitrocellulose, 0.45um, Whatman, Germany). After drying at room temperature for 15 minutes, the membranes were blocked with 5% of BSA in TBS containing 0.05% of Tween 20 (TBS-T), for 1 hour. The membranes were incubated with using anti-ApoA1 (1:1000, Santa Cruz, CA, USA), anti-ApoE and anti-ApoB (1:500, Santa Cruz, CA, USA) and anti human serum albumin (1:500, Santa Cruz, CA, USA). After washing with TBS-T, membranes were incubated with horseradish peroxidase-conjugated (Jackson ImmunoResearch, Tucker, GA, USA) secondary antibodies diluted 1:5000 in TBS-T with 1% BSA for 1 h.

For Western Blot analysis of EV markers, 5X Laemmli buffer was added to released EVs and sample boiled for 5 min at 95°C. Proteins were separated by SDS-PAGE (4– 20%, Mini-Protean TGX Precast protein gel, Bio-Rad) and transferred onto a nitrocellulose membrane (BioRad, Trans-Blot Turbo). Nonspecific sites were saturated with a TBS-T solution with 1% BSA for 1 h. Membranes were incubated overnight at 4°C with anti-CD9 (1:1000, BD Pharmingen, San Jose, CA, USA), anti-CD63 (1:1000; BD Pharmingen), anti-Alix (1:1000, Santa Cruz, CA, USA), and anti-TSG101 (1:1000, Novus Bio, Centennial, CO, USA). After washing with TBS-T, membranes were incubated with horseradish peroxidase-conjugated (Jackson ImmunoResearch, Tucker, GA, USA) secondary antibodies diluted 1:3000 in TBS-T with 1% BSA for 1 h.

For Western Blot analysis of RBC-EVs, 5X Laemmli buffer was added to released EVs and sample boiled for 5 min at 95°C. Proteins were separated by SDS-PAGE (10% polyacrylamide) and transferred onto a PVDF membrane. The blocking step was carried out with a PBS-T solution with 5% BSA for 1h at 37°C. Membranes were incubated overnight at 4°C with anti-CD9 (1:500, Santa Cruz, CA, USA), anti-CD81 (1:500, Santa Cruz, CA, USA) and anti-CD63 (1:1000, Merck-Millipore, MA, USA). Detection was achieved with secondary antibody anti-mouse HRP conjugate (Bethyl, TX, USA) and images were acquired with Chemibox Syngene).

### Transmission Electron Microscopy (TEM)

For negative staining 2uL pf sample was adsorbed on a glow discharged 300 mesh formvar/carbon coated grids and contrasted with 2% acqueous uranyl acetate solution. Grids were air dried and observe with a Talos L120C (FEI, Thermo Fisher Scientific) operating at 120kV. Images were acquired with a Ceta CCD camera (FEI, Thermo Fisher Scientific). For conventional TEM EVs absorbed on agarose magnetic beads were fixed with 2,5% glutaraldehyde in 0,1M cacodylate buffer. Using a magnetic stands the beads were washed in cacodylate buffer and postfixed with reduced osmium (1% OsO4, 1,5% potassium ferrocyanide in 0,1M cacodylate buffer pH 7.4) for 1 hour on ice. After several washes in milli-Q water samples were incubated in 0,5% uranyl acetate overnight at 4°C. Samples were then dehydrated with increasing concentration of ethanol, embedded in epoxy resin and polymerized in BEEM capsules for 48 hours at 60c°. Ultrathin sections (70-90 nm) were obtained using an ultramicrotome (UC7, Leica microsystem, Vienna, Austria), collected on copper or nickel grids, stained with uranyl acetate and Sato’s lead solutions and observed in a Transmission Electron Microscope Talos L120C (FEI, Thermo Fisher Scientific) operating at 120kV.

### Nanoparticle tracking analysis (NTA)

NTA was performed according to the manufacturer’s instructions using a NanoSight NS300 system (Malvern Technologies, Malvern, UK) configured with a 532 nm laser. Samples were diluted in micro-filtered PBS; the ideal measurement concentrations were identified by pre-testing the ideal particle per frame value (20–100 particles/frame). A syringe pump with constant flow injection was used and three videos of 60 s were captured and analyzed with Malvern NTA software version 3.4

### SiMoA beads conjugation to pan-teraspanin antibodies

Beads conjugation to antibodies was performed according to Quanterix Homebrew kit instructions using the recommended buffers as follows. Conjugation of 150 µl of carboxylate paramagnetic beads (2.8×10^9^ prt/ml) are washed three times with 300 µl of Bead Wash Buffer (Quanterix, phosphate buffer with detergent), after every washing step the beads are pulsed spin and placed on a magnetic separator for 1 minute to aspirate the supernatant. The beads are washed three more times with 300 µl of Bead Conjugation Buffer (Quanterix, 50 mM MES buffer pH6.2) and then are activated with EDC 0.3 mg/ml for 30 minutes at 4°C under mixing/shaking. 80 µg of antibody (CD9, CD63, CD81) are buffer exchanged with a 50 KDa Amicon filter and antibodies recovered in the Quanterix Bead Conjugation Buffer; after buffer exchange antibody concentration is measured with a Nanodrop spectrophotometer (ThermoFisher) and adjusted to 0.2 mg/ml with Bead Conjugation Buffer. 300 µl of a 0.2 mg/ml antibody solution are added to the activated paramagnetic beads and incubated for 2 hours at 4°C under mixing/shanking. After the conjugation step the beads are washed two times with Bead Wash Buffer and then are blocked with Bead Block Buffer (Quanterix, phosphate buffer with BSA) for 45 Minutes at room temperature under mixing/shaking. After blocking, beads are washed three times with Bead Diluent and stored until use at 4°C.

### SiMoA beads conjugation to MSP

150 µl of SiMoA carboxylate paramagnetic beads (2.8×10^9^ prt/ml) are activated with EDC according to Quanterix Homebrew kit instructions as described above, then 300uL NH2-Maleimide linker (from Sigma-Aldrich) solution 10 mM in PBS (adjusted to pH 8.6) is added and shaked for 2 hours in RotoFlex. Beads are then washed 2 times with PBS to remove NH2-Maleimide in excess and incubated with 300uL of 100uM solution of MSP in PBS (adjusted to pH 8.6, with 2 equivalents DIEA and 1 mM TCEP). Peptide reacts for 1 hour under mixing. After the conjugation step the beads are washed two times with PBS and then are blocked with Bead Block Buffer (Quanterix) for 15 Minutes at room temperature under mixing/shaking. After blocking, beads are washed with Bead Wash Buffer (Quanterix) and stored in Bead Diluent (Quanterix) at 4°C.

### Pan-tetraspanin three-step assay

Pan-tetraspanin beads solution are prepared at the concentration of 2×10^7^ beads/ml in Bead Diluent. The detector antibody (biotinylated CD9, CD63, CD81 antibodies by Ancell or anti-band 3 from Santa Cruz) solutions (0.3 µg/ml) are diluted in Homebrew Sample Diluent (Quanterix); similarly, serum samples are diluted 1:4 in Homebrew Sample Diluent (Quanterix) whereas plasma samples are diluted 1:10 in Homebrew Sample Diluent. 25 µl of beads are transferred into a 96 microwell plate and mixed with 100 µl diluted sample and incubated for 30 minutes at 25°C at 800rpm. After incubation, beads are washed with an automatic plate-washer and then incubated for 10 minutes with 100 µl of detector antibody After incubation, beads are washed and incubated for 10 minutes with a 150 pM SBG solution (in SBG Diluent, Quanterix). After SBG incubation step the plate is washed again and then inserted into the Quanterix SR-X instrument for analysis where RGP is automatically added. Data were analyzed and processed by Reader Software Simoa® 1.1.0.

### MSP SiMoA three-step assay

The assay is run as described above for pan-tetraspanin beads except that samples and detector antibodies are incubated in PBS. The detector antibody (biotinylated CD9, CD63, CD81 antibodies or or CD42a and CD62P by MiltenyBiotech or anti-band 3 from Santa Cruz) is used at the concentration of 0.6 µg/ml, serum samples are diluted 1:4, plasma samples are diluted 1:10. 25 µl of beads are transferred into a 96 microwell plate and mixed with 100 µl diluted sample and incubated for 30 minutes at 25°C at 800rpm. After incubation, beads are washed with an automatic plate-washer using optimized Tween concentration and then incubated for 10 minutes with 100 µl of detector antibody. After that, beads are washed with an automatic plate-washer and incubated for 10 minutes with a 150 pM SBG solution (in SBG Diluent, Quanterix). After SBG incubation step the plate is washed and then inserted into the Quanterix SR-X instrument for analysis where RGP is automatically added. Data were analyzed and processed by Reader Software Simoa® 1.1.0.

### RBC EV

RBCs obtained from anonymized type 0+ healthy volunteers under written consent were provided by the blood transfusion unit of A. O. Spedali Civili di Brescia (ethical approval NP5705) in sealed sterile bags. RBCs EVs were isolated using Ca^2+^/Ca^2+^ ionophore induction, following the guidelines from Usman et al. Briefly, RBCs were pelleted by centrifugation at 1,000×g for 8 minutes at 4 °C, and washed thrice in sterile PBS w/o Ca^2+^ and Mg^2+^. RBCs were further washed twice with CPBS (sterile PBS + 0.1 g L^−1^ CaCl) and transferred into 175 mm^2^ tissue culture flasks. Calcium ionophore (A23187, Sigma-Aldrich) was added to the flasks (final concentration 10 mM) and incubated overnight at 37 °C. RBCs were gently collected from the flasks, and cellular debris were removed by differential centrifugation (600×g for 20 min, 1,600×g for 15 min, 3,260×g for 15 min, and 10,000×g for 30 min at 4 °C), discarding the pellet after each centrifugation step and transferring the supernatant into fresh sterile tubes. The supernatants were filtered through 0.45 μm nylon syringe filters (Nalgene). EVs were collected by ultracentrifugation at 50,000×g for 70 min at 4 °C. The pellets were then resuspended in cold sterile PBS, layered above a 2 mL frozen 60% sucrose cushion, and centrifuged at 50,000×g for 16 h at 4 °C, with the deceleration speed set to 0. The red layer of EVs was collected and washed twice with cold sterile PBS and spun at 50,000×g for 70 min at 4 °C. Finally, EVs were resuspended in 1 mL of cold sterile PBS, aliquoted and stored at −80 °C until used.

Centrifugations below 10,000xg were performed on an Eppendorf 5804 R equipped with a A-4-44 swinging bucket rotor. Ten thousand x g step was performed on a Beckman Avanti centrifuge equipped with a JA-20 fixed angle rotor. A Beckman XPN-80 equipped with a TY45-Ti fixed angle rotor was employed for the ultracentrifugation step. Sucrose cushion ultracentrifugation was performed on a Beckman Optima Max-XP equipped with a MLS-50 swinging arms rotor. The final washing step was performed om a Optima MAX-XP equipped with a TLA-55 rotor.

### Serum and plasma samples for the clinical validation

Peripheral venous blood samples were collected from patients recruited at the Istituto Cardiocentro Ticino, Ente Ospedaliero Cantonale (Lugano, Switzerland). The study protocol was approved by the local ethical committees. All participants gave informed written consent to the study in accordance with the declaration of Helsinki. Peripheral venous blood samples were collected from patients presenting with a diagnosis of STEMI, according to the European Society of Cardiology (ESC) guidelines ^51^ on presentation to the emergency department before primary PCI. In addition, samples were collected from patients with chronic CAD presenting with stable angina (SA) according to ESC guidelines ^51^ and age-matched healthy control subjects. For serum blood was collected in heparin-free polypropylene tubes, while for plasma (only in STEMI patients) in sodium citrate tubes, and centrifuged at 1,600g for 15 minutes at 4°C degree to separate and discard cellular components. Serum, and free-platelet plasma were then differentially centrifuged at 3,000g for 20 min, at 10,000g for 30 min, and at 20,000g for 15 min as previously described ^47^; supernatant was aliquoted, stored at −80°C, and never thawed prior to analysis.

### Statistical analysis

Statistics was performed by IBM SPSS Statistics 25 (Armonk; NY) and GraphPad PRISM 9.0 (La Jolla, California). EV marker expression was compared by Kruskal-Wallis and Mann-Whitney tests. Correlations of expression levels in serum and plasma were assessed by Pearson’s R test and analysis of the regression curves. The analysis of Bland-Altman plots was used to assess the within-sample relationship and detect systematic, proportional, or magnitude-dependent biases. A P-value lower than 0.05 was considered significant. The analysis of receiver operating character-istics (ROC) curves was used to compare diagnostic performances of selected variables. Multivariate logistic regression analysis was performed to determine odds ratio (ORs). P-values <.05 were considered significant

