## Supplementary material for "Addressing Heterogeneity in direct analysis of Extracellular Vesicles and analogues using Membrane-Sensing Peptides as Pan-Affinity Probes": Supp Figures

*1: Consiglio Nazionale delle Ricerche, Istituto di Scienze e Tecnologie Chimiche "Giulio Natta" (SCITEC), Milano, Italy*

*2: Cardiovascular Theranostics, Istituto Cardiocentro Ticino, Ente Ospedaliero Cantonale, CH-6500, Bellinzona, Switzerland*

*3: Institute for Research in Biomedicine, Faculty of Biomedical Sciences, Università della Svizzera italiana (USI), CH-6500, Bellinzona, Switzerland*

*4: Department of Molecular and Translational Medicine, University of Brescia, Viale Europa 11, 25123 Brescia, Italy*

*5: CSGI, Center for Colloid and Surface Science, 50019 Florence, Italy*

*6: Euler Institute, Faculty of Biomedical Sciences, Università della Svizzera italiana, 6900 Lugano, Switzerland*

§ : Equally contributed

*: Corresponding authors

### Supplementary Information

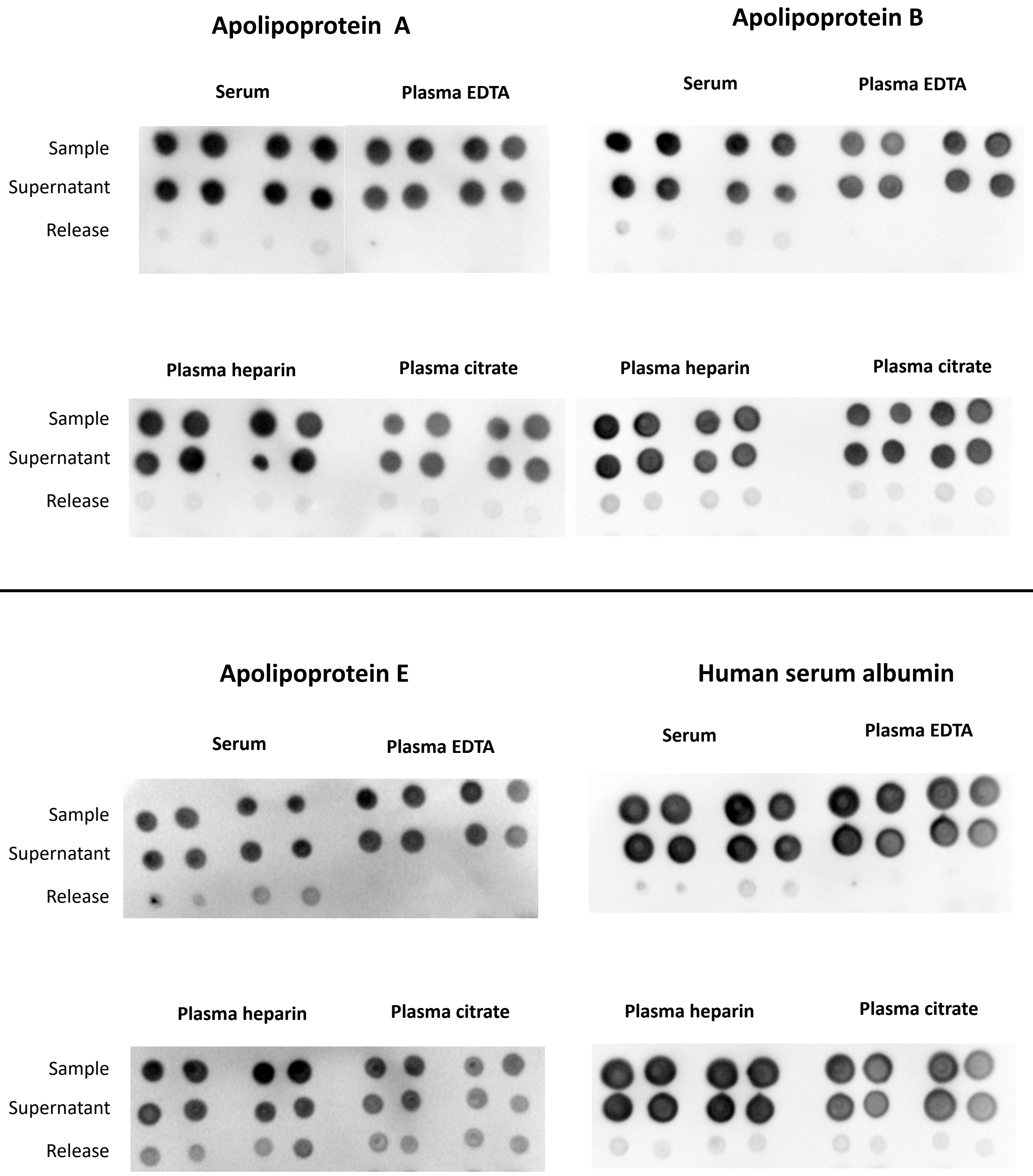


**Figure S1:** Immune dot-blot for the evaluation of Alipopoprotein A, Apolipoprotein B, Apolipoprotein E and serum albumin in starting sample, supernatant, release fraction. Blood was collected from healthy subjects, and four preanalytical conditions evaluated: serum, Plasma EDTA, Plasma Citrate, Plasma heparin. Plasma and serum were isolated in parallel from the same subject. EDTA, heparin and citrate tubes were used for the collection of plasma, while serum was obtained in clot activator tubes. Two centrifuge steps were performed for all samples: 1500g for 10 minutes and 2500g for 10 minutes.

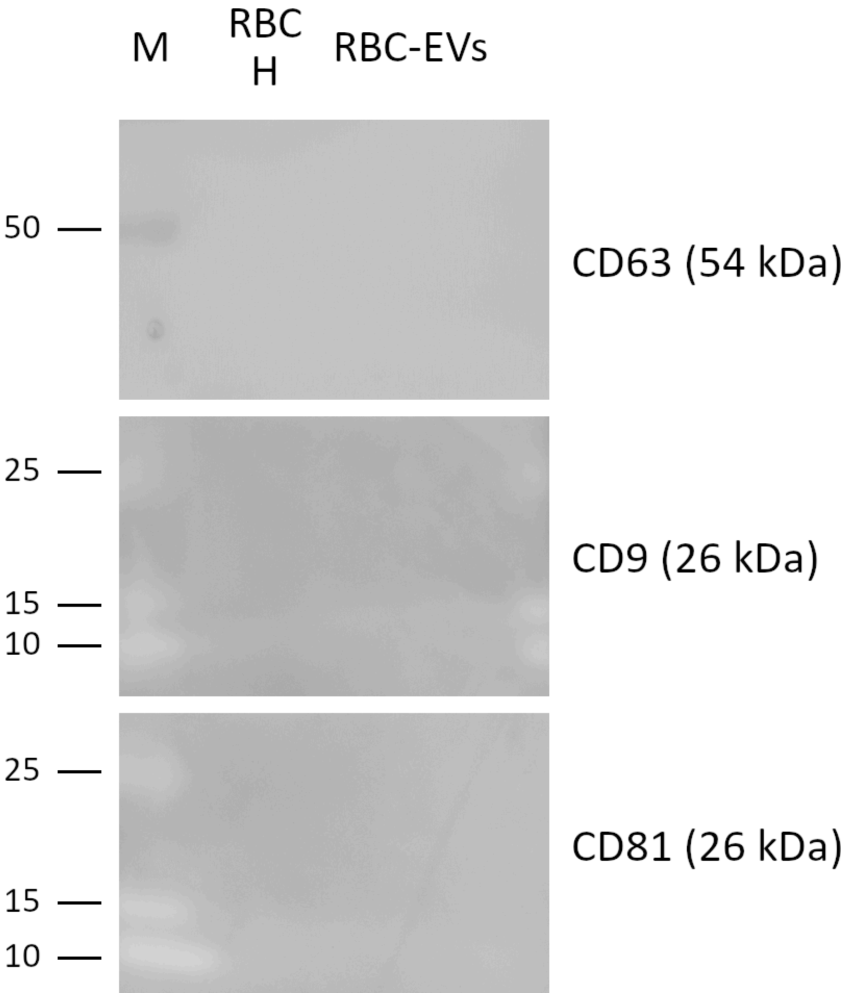


**Figure S-2:** RBC-EVs poorly express the exosomal tetraspanin triad CD63, CD9, and CD81, reflecting their biogenesis pathway (budding from RBC plasma membrane). Legend: M = Marker; RBC-H = RBC homogenate; RBC-EVs = RBC EVs ectosomes.
